# Copper-Induced Stress and Recovery Impacts on Organismal Phenotypes and the Underlying Proteomic Signatures in *Botryllus schlosseri*

**DOI:** 10.1101/2025.06.12.659394

**Authors:** Maxime Leprêtre, Dietmar Kültz

## Abstract

This study establishes the copper tolerance range of the colonial marine tunicate *Botryllus schlosseri*. Furthermore, quantitative organismal phenotyping and quantitative proteomics were combined to characterize the *B. schlosseri* response to, and recovery from, acute copper exposure stress. Changes in the area of *B. schlosseri* colony systems and pigmentation provided sensitive, dose-dependent markers of exposure to, and recovery from, copper stress. Comprehensive quantitative proteomics using consistent data-independent acquisition (DIA) assay libraries revealed activation of detoxification, oxidative stress, and immune pathways during exposure to copper stress. In addition, quantitative proteomics uncovered enrichment of tissue remodeling and proliferative signaling pathways during recovery from copper stress. We identified 35 proteins whose expression closely mirrored phenotypic changes observed at the colonial system level. This specific proteome signature represents a comprehensive molecular underpinning of the organismal response of *B. schlosseri* to copper stress. In conclusion, this study establishes copper tolerance ranges of the invasive colonial tunicate *B. schlosseri* and explains the molecular underpinnings of stress-induced organismal phenotypes by identifying corresponding proteome signatures and their associated functional enrichments. Moreover, identification of copper concentrations that are stressful and highly disruptive on the molecular phenotype, yet readily recoverable from, lays a critical foundation for future studies directed at stress-induced adaptation and evolutionary trajectories of marine invertebrates in changing and novel environments.

## 1. Introduction

Marine organisms regularly face stressful environmental changes that affect their physiology, fitness, and survival. Among these stressors, anthropogenic pollutants, particularly heavy metals, have become ubiquitous in coastal ecosystems (Shah, 2021). Copper, one of the most pervasive heavy metals, is introduced into marine environments through human activities, including the use of antifouling paints, mining operations, and industrial discharges (Rehman et al., 2019). While copper is an essential micro-nutrient in trace amounts, continuous exposure to copper can be toxic to organisms if their tolerance limits are exceeded (Rehman et al., 2019). This is specifically true for marine invertebrates where excess of copper has been shown to cause oxidative stress, cellular damage, immune dysfunction and, in extreme cases, death (Feng et al., 2022; Li et al., 2023). Moreover, owing to accelerating industrialization and global environmental changes, copper toxicity is expected to intensify. Such continuously shifting conditions may amplify the harmful effects of this pollutant and the adaptive advantages of invasive species over long-standing stenotopic local species (Cui et al., 2024).

In addition to its acute toxicity, copper acts as a strong agent of natural selection, prompting rapid evolutionary adaptations in various aquatic species (Bourret et al., 2008; Hochmuth et al., 2015; Han et al., 2025). This adaptive process represents an illustration of the broader concept known as stress-induced evolution (SIE). The SIE concept states that environmental stressors accelerate the emergence of heritable phenotypic variation through mechanisms such as increased mutation rates and epigenetic modifications (Mojica and Kültz, 2022). In ascidians, for example, increased tolerance to copper has been linked to successful invasion of, and adaptation to, novel environments (Osborne et al., 2018). Such invasion-facilitating increase in copper tolerance illustrates how copper-induced stress can shape population dynamics, resilience, and distribution ranges. These findings identify copper as a relevant model stressor for investigating the role of environmental stress on the swiftness of evolutionary processes in marine organisms.

Within this context, the highly invasive colonial tunicate *Botryllus schlosseri*, and its copper tolerance, stands out as a particularly intriguing model for investigating the physiological and molecular foundations of SIE. *B. schlosseri* forms colonies of genetically identical zooids organized into functional units called systems that are interconnected by a common vascular network (Manni et al., 2019). Its asexual reproduction via weekly budding (blastogenesis) enables continuous regeneration and rapid colony growth (Gasparini et al., 2015; Manni et al., 2019). This clonal and highly regenerative biology allows for repeated, synchronized sampling of genetically uniform individuals across time and conditions, minimizing genetic variability and enabling precise comparisons of stress responses and stress-induced changes. In addition to its unique developmental biology, *B. schlosseri* exhibits high tolerance to environmental fluctuations and a strong invasive potential (Cima et al., 2015; Tobias et al., 2024), traits that underscore its ecological plasticity and high adaptive capacity. These features render it especially relevant for studying how (and how rapidly) environmental stressors, such as copper pollution, drive physiological responses, evolutionary change, and invasion of novel environments in marine invertebrates.

Despite this strong potential, our understanding of *B. schlosseri* stress responses remains limited. Most existing studies rely on qualitative morphological markers, such as ampullae condition and other visible traits to assess stress in colonial tunicates (Gregorin et al., 2021; Rosner and Rinkevich, 2024). While informative, these approaches lack the sensitivity and objectivity needed to detect early or subtle physiological changes that could be instrumental in facilitating adaptation to novel or altered environments. Furthermore, molecular signatures associated with *B. schlosseri* responses to specific types of stress remain largely unexplored, with the exception of immune-related genes (Rosner and Rinkevich, 2024). This is particularly true for the proteome level, which represents the closest molecular proxy of higher-order, organismal phenotypes. The lack of quantitative phenotypic signatures and comprehensive information on molecular phenotypes limits our ability to understand how *B. schlosseri* copes with, adapts to, and evolves under environmental stress such as high copper exposure.

To address these knowledge gaps, the present study combines quantitative organismal phenotyping with data-independent acquisition (DIA) proteomics. In addition, this is only the second quantitative proteomics study performed on any colonial tunicate, following up on a previous comparison of proteomes from field and laboratory *B. schlosseri* populations (Kültz et al., 2024). Combining the analysis of molecular and organismal phenotypes permits comprehensive assessment of *B. schlosseri* responses to different levels of copper exposure, as well as in-depth mechanistic understanding of their capacity to recover once the stressor is removed. The objectives of this study were to (i) identify robust phenotypic indicators and molecular signatures of copper stress, (ii) characterize the key biological pathways involved in both the response to, and recovery from, copper stress, and (iii) generate foundational data for experimentally probing stress-induced evolution (SIE) in longer-term experiments that utilize periodical exposure to, and recovery from, most severe but non-lethal copper stress that greatly alters the *B. schlosseri* phenotype subject to natural selection. By capturing both immediate and reversible responses, this study provides a framework for future investigations into long-term adaptation and epigenetic/ evolutionary dynamics. Ultimately, this work contributes to a broader understanding of how marine organisms respond to and recover from environmental stressors in the face of increasing anthropogenic (e.g. industrial and urban pollution), as well as natural (e.g. volcanism) pressures.

## 2. Material and method

### 2.1. Sampling and maintenance of animals

Two wild *B. schlosseri* genotypes (G1 and G2) were collected from floating docks in Berkeley Marina. Larvae from these colonies settled on glass slides within one week of spawning at the UC Davis Cole B facility. These lab-born colonies were reared in a recirculating aquaculture system (RAS; salinity: 30ppt, temperature: 19°C, pH: 8) and allowed to attach vertically to the glass. A third genotype (G3) was likely produced through sexual reproduction within the laboratory’s RAS or may have originated from other field collection, though its exact origin is uncertain.

Each genotype was kept isolated in separate tanks to prevent inter-colony competition and cross-fertilization. Colonies were fed daily a mixture of live algae (*Dunaliella salina, Tetraselmis chuii, Isochrysis galbana*, and *Nannochloropsis oculata*) complemented with commercial rotifer-based liquid feed (Roti-Rich, Florida Aqua Farms). Colonies were gently cleaned weekly using soft brushes. All colonies were maintained in stable RAS conditions for at least three months before experiments.

### 2.2. Copper Exposure and Recovery of Botryllus schlosseri Colonies

To assess the stress response of *B. schlosseri*, systems from the three genotypes (G1, G2, and G3) were acclimated for two weeks in 2-liter glass jars filled with recirculating aquaculture system (RAS) seawater. During acclimation, tunicates were maintained under a 12-hour light/dark cycle and fed ad libitum with a mixture of live microalgae.

To standardize physiological conditions, copper exposures were initiated 24 hours after the takeover (TO) stage of the blastogenic cycle. *B. schlosseri* systems were transferred to 600 mL plastic beakers containing seawater from the RAS. Stock solutions of copper were prepared using copper sulfate pentahydrate (CuSO ·5H O). Nominal copper concentrations of 0 µg/L (C0), 30 µg/L (C1), 100 µg/L (C2), 300 µg/L (C3), and 900 µg/L (C4) were achieved by adding 150 µL of the appropriate stock solution to 600 mL of seawater. To avoid potential confounding effects, colonies were not fed during the 24-hour copper exposure period. After exposure, colonies were returned to 2-liter jars without copper and maintained under the same acclimation conditions (light/dark cycle, temperature, and feeding regime) until the completion of their blastogenic cycle.

For each copper concentration and genotype, a minimum of five systems (each comprised of at least six zooids) were analyzed. The experiment was repeated three times, resulting in at least 15 biological replicates per genotype and treatment. Morphological changes were observed daily using a stereomicroscope. Images were captured at predefined time points: baseline (BL), 24 hours post-exposure (E24), and 24 (R24), 48 (R48), and 96 (R96) hours post-exposure recovery.

### 2.3. Monitoring Size and Pigmentation Changes in Botryllus schlosseri Systems

Morphological and pigmentation changes were monitored using stereomicroscopic images acquired at consistent magnification and illumination settings. Backgrounds, including tunics, were manually removed in Microsoft PowerPoint to isolate individual systems composed of grouped zooids and buds. The cleaned images were then saved in JPG format for subsequent analysis.

Image analysis was conducted using Fiji (ImageJ). Images were converted to 8-bit grayscale, and tunicate systems were segmented using a standardized threshold to minimize background artifacts. Quantitative data were extracted from each image, including total area and mean grayscale intensity, which served as proxies for pigmentation and area of systems over time. For each system, area and grayscale intensity values were measured at E24, R24, R48, and R96, and expressed as a percentage change relative to the baseline (BL). The non-parametric Kruskal– Wallis test was used to assess overall differences in area and pigmentation changes across exposure conditions. When significant, post-hoc pairwise comparisons were conducted using Dunn’s test with Bonferroni correction. Differences were considered statistically significant at an adjusted p-value < 0.05.

### 2.4. Proteomic investigations

#### 2.4.1. Experimental conditions for Proteomic Analysis

Quantitative proteomics was conducted exclusively on genotype G1. Copper concentrations of 300 µg/L (C3) and 900 µg/L (C4) were chosen based on phenotypic responses observed in prior assays. As previously described, colonies were exposed for 24 hours (E24) to three copper concentrations (C): 0 µg/L (C0, control), 300 µg/L (C3), and 900 µg/L (C4). Each treatment included 11 to 13 systems, with each system comprised of at least six zooids.

To examine recovery processes, approximately half of the colonies from each exposure condition were transferred to clean RAS seawater for an additional 24 hours (R24). For both exposure and recovery conditions, we collected either individual large systems or pooled smaller ones to obtain a minimum of 15 zooids per biological replicate. Dissections included not only the zooids within systems but also the surrounding vascular network and peripheral ampullae present under a single tunic.

The experiment was independently repeated three times, yielding a total of 11 biological replicates per condition. Samples were flash-frozen in liquid nitrogen immediately after exposure or recovery and stored at −80 °C until protein extraction.

#### 2.4.2. Sample Preparation for Mass Spectrometry

Samples underwent chemical and mechanical lysis to optimize protein extraction. Each sample was lysed in 8 M urea and 50 mM ammonium bicarbonate (AmBic), pH 7.0. To facilitate homogenization, approximately 20 zirconium beads (1 mm diameter; Benchmark Scientific, Cat. No. D1031-10) were added and samples were processed using a BeadBug™ microtube homogenizer (Benchmark Scientific) at 40,000 rpm for 30 seconds. Lysates were centrifuged at 12,000 × g for 10 minutes, and the resulting supernatants were transferred to 1.5 mL low-binding Eppendorf tubes. Proteins were precipitated by adding cold acetone with 10% trichloroacetic acid (TCA) to achieve a 5:1 ratio (acetone:lysate, v/v), followed by incubation at −20°C for 1 hour. The samples were centrifuged at 15,000 × g for 10 minutes, and the supernatant was discarded. Pellets were washed twice with cold acetone, centrifuged again at 15,000 × g for 10 minutes, air-dried, and resuspended in 50 µL of lysis buffer. Reduction was performed by adding dithiothreitol (DTT) to a final concentration of 5 mM, followed by incubation at 60°C for 10 minutes. Alkylation was achieved by adding iodoacetamide (IAA) to 15 mM and incubating at room temperature in the dark for 30 minutes. Residual IAA was quenched with 5 mM DTT. Protein concentration was determined using the Pierce™ 660 nm Protein Assay (Thermo Fisher Scientific, Cat. No. 22660).

For digestion, urea was diluted to below 1.5 M using LC-MS grade water and 1 M AmBic (pH 8.5), maintaining a final AmBic concentration of 50 mM. Proteins were digested using Pierce™ Trypsin/Lys-C Protease Mix, MS Grade (Thermo Fisher Scientific, Cat. No. A40007) at a 1:50 enzyme-to-substrate ratio (w/w), based on estimated protein content. Digestion proceeded at 37°C for 3 hours with shaking at 250 rpm. The reaction was stopped by adding formic acid to a final concentration of 2%. The peptides were desalted using Pierce™ C18 Spin Columns (Thermo Fisher Scientific, Cat. No. 89870) according to the manufacturer’s instructions. Elution was performed with 50% acetonitrile, followed by drying in a speed vacuum and resuspension in 0.1% formic acid. Final peptide concentrations were determined using the Pierce™ Quantitative Fluorometric Peptide Assay (Thermo Fisher Scientific, Cat. No. 23290) and normalized to 100 ng/µL.

#### 2.4.3. Mass spectrometry analysis

Liquid chromatography–mass spectrometry (LC-MS) analysis was performed using a Bruker nanoElute® 2 system coupled to a ultra high-resolution (UHR) QTOF mass spectrometer (Impact II, Bruker) operating in positive ion mode. A total of 100 ng of peptides per sample was injected and separated on an Aurora Ultimate CSI C18 reverse-phase analytical column (25cm x 75µm x 1.7µm, Ion Opticks) maintained at 50°C. Peptides were eluted over a 50-minute linear gradient from 3% to 33% acetonitrile in 0.1% formic acid, at a flow rate of 300 nL/min. Mass spectrometric data were acquired using a data-independent acquisition (DIA) mode with 42 isolation with windows, each covering a 20 m/z mass range with 0.5 m/z overlap, and 0.86s cycle time.

Raw DIA data files were converted to mzML format using MSConvert (ProteoWizard). Precursor identification and quantification were carried out using DIA-NN (version 2.1.0). A spectral library was generated in silico from the predicted proteome of *B. schlosseri*, based on the genome published by Thier et al. (2024). Trypsin was specified as the proteolytic enzyme, allowing up to two missed cleavages. Carbamidomethylation of cysteine was set as a fixed modification, while methionine oxidation and N-terminal acetylation were included as variable modifications. This predicted spectral library was then used for precursor identification and quantification in DIA-NN, which was run with default parameters.

#### 2.4.4. Comparative proteomic analysis

To assess changes in protein abundance across experimental conditions, data were normalized and quantified using the DirectLFQ algorithm (Ammar et al., 2023). Protein groups supported by at least two unique peptides were retained for downstream analysis. In addition, protein groups with more than 50% missing values across the entire dataset were excluded to ensure data quality and reliability.

Statistical analyses were performed using the ProLFQua R-package (Wolski et al., 2023), applying linear models with empirical Bayes moderation to enhance sensitivity. The Benjamini-Hochberg procedure was used to control the false discovery rate (FDR). To evaluate proteomic responses to copper exposure, protein abundances in the C3 and C4 treatment groups were compared to the control condition (C0). To identify proteins involved in recovery, comparisons were made between exposure (E24) and recovery (R24) time points within the same treatment groups (C0, C3 and C4). Proteins with an FDR < 0.05 were considered differentially abundant (DAPs). Venn diagrams and UpSet plots were used to visualize DAPs shared or unique across experimental conditions. They were created with the VennDiagram and ComplexUpset R-packages, respectively.

K-means clustering analysis of DAPs was performed using the R-package ClusterGVis. Prior to clustering, the intensities of differentially abundant proteins (DAPs) were log transformed and standardized (Z-score normalization) across samples. To simplify the analysis and highlight condition-specific trends, protein abundances were averaged across replicates within each experimental condition. The optimal number of clusters (K) was determined by evaluating the within-cluster sum of squares across a range of K values. Once the optimal K was selected, K-means clustering was applied to the averaged DAP data. Cluster assignments were visualized using line plots and heatmaps, with rows representing individual DAPs and columns corresponding to the experimental conditions. Additional heatmaps and boxplots were generated using the ComplexHeatmap and ggplot2 packages in R.

#### 2.4.5. Functional analyses

The functional annotation of protein sequences was conducted using a dual-methodology approach. Initially, DIAMOND was utilized to perform BLASTP searches against the SwissProt database without any taxonomic limitations, retaining only annotations with E-values below 1e-3 In parallel, the eggNOG-mapper tool was employed to identify orthologous groups and extract KEGG (Kyoto Encyclopedia of Genes and Genomes) identifications and pathway annotations (Kanehisa and Goto, 2000; Cantalapiedra et al., 2021).

Functional enrichment analyses were conducted using the clusterProfiler R-package (Wu et al., 2021) to investigate the biological functions linked to DAPs related to copper exposure and recovery. Over-representation analyses were performed using a custom annotation database derived from eggNOG. The background set comprised all protein groups identified in the proteomic dataset, while the test sets included DAPs specific to individual conditions or grouped by k-means clustering. KEGG pathways with a p-value below 0.05 were considered significantly enriched.

## 3. Results

### 3.1. Morphological and Physiological Changes During Copper Exposure and Recovery

This study explores the phenotypic responses of *B. schlosseri* to copper exposure, aiming to identify reliable bioindicators for environmental stress assessment. Among the earliest and most obvious morphological alterations following copper exposure were changes in system pigmentation and area in all three *B. schlosseri* genotypes (Figure 1, Supplementary Figure S1). Under non-exposed conditions, *B. schlosseri* systems typically became lighter in color and increased in area during the course of blastogenesis. In contrast, copper-exposed systems became progressively darker and smaller (Figure 1, Supplementary Figure S1).

**Figure 1.**
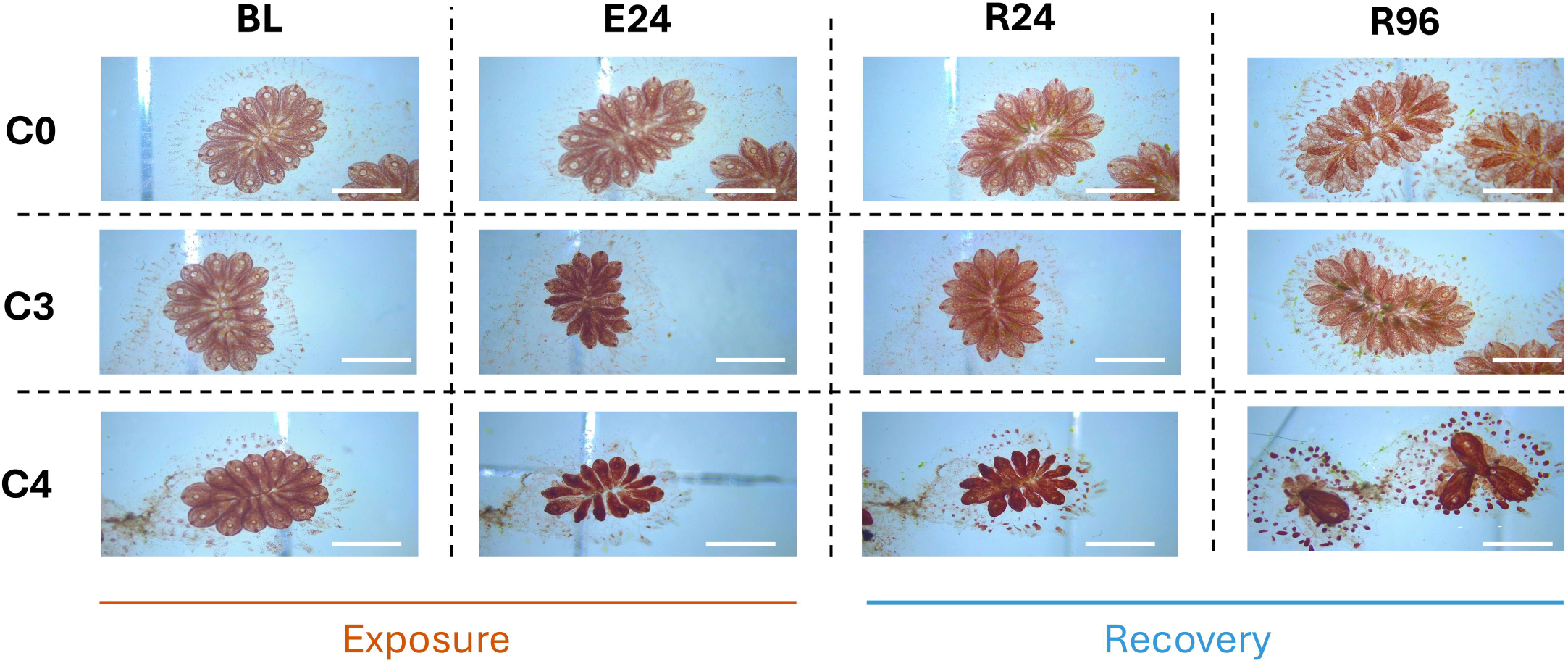
**Visual Comparison of Tunicate Responses to Copper Exposure and Recovery**. Representative images of genotype 1 (G1) tunicate systems at baseline (BL), after 24Lh of copper exposure (E24), and during recovery at 24Lh (R24) and 96Lh (R96), at three copper concentrations: C0 (0Lµg/L), C3 (300Lµg/L), and C4 (900Lµg/L). All images were taken at the same magnification and light intensity to allow direct visual comparison.

#### 3.1.1. Dose-Dependent Effects on System Area

A clear dose-dependent reduction in system area was observed 24 hours after copper exposure (E24), with increasing concentration leading to progressively greater shrinkage (Figure 2A). In the control condition (C0), zooids exhibited about 25% increase in area at E24 relative to baseline (BL), consistent with expected growth during blastogenesis. Colonies subjected to low copper concentrations (C1 and C2) showed system areas similar to those of the control group. However, a slight reduction in zooid area was observed at C2 compared to C1, which was only significant in genotype G1, indicating early signs of sensitivity to copper at moderate concentrations. At higher concentrations (C3 and C4), system area reduction became more pronounced and was significantly different from lower exposure groups (C0, C1, and C2). Specifically, systems exposed to 300 µg/L (C3) showed area reductions exceeding 25%, while those exposed to 900 µg/L (C5) exhibited area reductions of over 50%, highlighting a strong dose-dependent impairment of *B. schlosseri* structural integrity.

**Figure 2.**
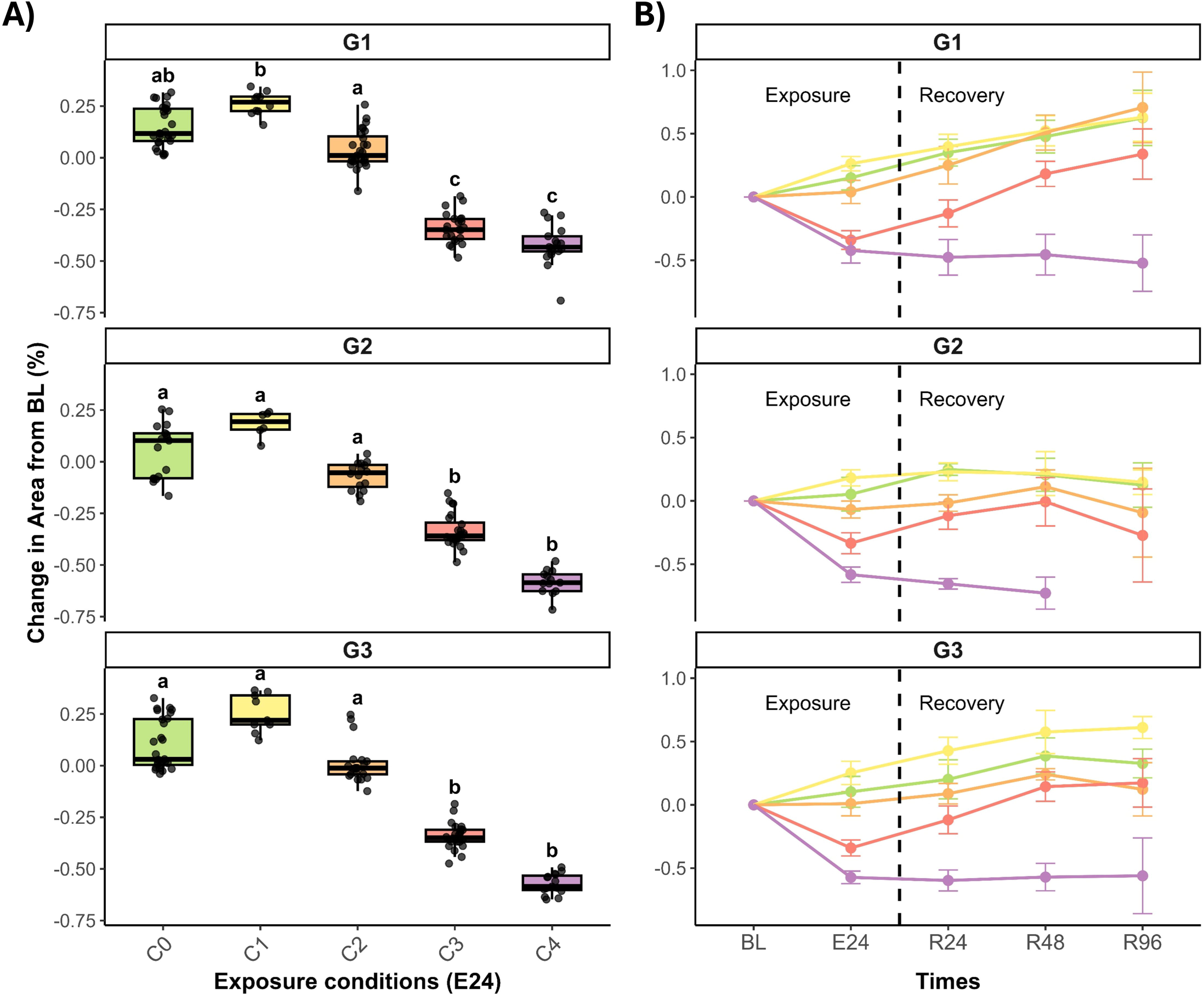
Dose-dependent effects of copper exposure on system area and recovery across the three genotypes. (A) Boxplots showing the percentage change in system areas relative to baseline (BL) after 24 hours of exposure (E24) across five copper concentrations (C0 = 0 µg/L, C1 = 30 µg/L, C2 = 100 µg/L, C3 = 300 µg/L, C4 = 900 µg/L) in genotypes G1, G2, and G3. Different letters indicate statistically significant differences between conditions within each genotype (Tukey’s HSD, p < 0.05). (B) Line plots showing the dynamics of system area change over time, during exposure (BL–E24) and recovery (R24–R96) for each genotype. Line colors correspond to copper concentrations and points represent mean values ± standard error. Dashed vertical line marks the end of exposure and beginning of the recovery.

#### 3.1.2. Recovery Dynamics After Copper Exposure

Upon recovery, colonies exhibited divergent responses depending on the level of prior copper exposure (Figure 2B). In C0, C1, and C2 treatments, system areas gradually increased over time, reaching up to a 50% expansion by the end of the blastogenic cycle (R96), reflecting normal growth patterns. Systems exposed to 300 µg/L (C3) demonstrated partial recovery where system areas began expanding within 24 hours and continued to increase, nearing control levels by R96. In contrast, zooids exposed to 900 µg/L (C4) failed to recover by R96. Shrinkage persisted and even worsened over time, indicating that high copper concentrations severely compromise the regenerative capacity of the colonies. By the fourth day of recovery (R96), many zooids in the C4 group had undergone reabsorption, while others developed multiple buds in genotypes G1 and G3, suggesting that budding continued despite the severe stress (Figure 1, Supplementary Figure S1). However, in genotype G2, all zooids exposed to the C4 concentration died before completing their blastogenic cycle, characterized by a distinctive red pigmentation and halted hemolymph circulation (Supplementary Figure S1A).

#### 3.1.3. Changes in Pigmentation During Exposure and Recovery

Parallel to changes in area, system pigmentation followed a dose-dependent pattern during the exposure phase (Supplementary Figure S2). In the control and low-exposure groups (C0, C1, and C2), pigmentation remained consistent with baseline (BL) levels after 24-hours exposure (Supplementary figure S2A). However, at higher copper concentrations, pigmentation intensified significantly. Systems exposed to 300 µg/L (C3) showed noticeable darkening after exposure, while those exposed to 900 µg/L (C4) exhibited even more pronounced pigmentation changes, reflecting a strong phenotypic response to copper exposure at these higher concentrations. During the recovery phase, pigmentation changes further revealed the contrasting results between moderate and high copper exposure (Supplementary Figure S2B). Systems from the treatment C3 gradually recovered their original coloration and approached control pigmentation levels at R96. Conversely, systems exposed to the highest concentration of copper (C4) continued to darken throughout the recovery period, indicating sustained stress impacts and cellular deterioration at R96.

#### 3.1.4. Additional Physiological Effects and Blastogenic Cycle Alterations

In addition to system shrinkage and pigmentation changes, copper exposure also induced pigmentation in the vasculature of colonies, led to emptying of the digestive system, and triggered irregular and involuntary zooid contractions. These changes highlight further physiological consequences of copper-induced stress. Copper also disrupted the timing of the blastogenic cycle. In genotypes G1 and G2, the blastogenic cycle was delayed by more than 12 hours at concentration C2 compared to C0. This delay increased proportionally to copper concentration, reaching over 24 hours at 300 µg/L (C3) in genotype G2. Interestingly, at the highest concentration (C4), the duration of the blastogenic cycle was comparable to the control. Despite signs of stress such as reabsorption of zooids, budding continued in colonies exposed to 900 ug/L of copper (C4). This divergence between reabsorption and budding suggests a shift in energy allocation under extreme stress, possibly prioritizing budding over zooid recovery.

### 3.2. Proteomic Signatures of Copper Stress and Recovery

Based on the observed phenotypic changes, a proteomic analysis was conducted to investigate the molecular mechanisms underlying the response to copper exposure and subsequent recovery. To this end, DIA investigations were performed on colonies of the G1 genotype following 24 hours of copper exposure (E) and after 24 hours of recovery (R), at copper concentrations of 0, 300, and 900 µg/L. A total of 66 samples were analyzed using DIA LC-MS, resulting in the identification of 7,025 protein groups and 43,475 related peptides. After filtering out proteins detected with less than 2 peptide sequences and not detected in at least 50% of the samples, a final set of 4,072 protein groups was retained for comparative proteomic analysis.

#### 3.2.1. Proteomic response to copper exposures

Comparative quantitative proteomics of *B. schlosseri* exposed to copper revealed a clear dose-dependent change in molecular phenotypes (proteomes) underlying the organismal phenotype changes outlined above. In tunicates exposed to 300 ug/L of copper (C3), 223 DAPs were identified compared to control conditions (C0). This number almost tripled at C4 (582 DAPs) indicating a broader and more intense molecular response at a higher exposure level of copper. Most proteins affected at C3 (193 DAPs) were also altered at C4, indicating utilization of common molecular pathways, with stronger effects at higher copper concentration. K-means clustering analysis of these DAPs was performed to explore their dynamic patterns across different copper concentrations, revealing four distinct expression clusters (Figure 3). Clusters 1 and 2 included 142 and 169 DAPs, respectively, that increased in abundance compared to control conditions. In Cluster 1, the increase was greater in C4 than in C3, whereas Cluster 2 contained DAPs that were already elevated in C3 and remained high or showed a slight further increase in C4. A similar pattern was observed for downregulated DAPs where Cluster 3 (127 DAPs) showed a moderate further decrease in C4 compared to C3, while Cluster 4 (170 DAPs) exhibited a more pronounced abundance decrease in C4. Among the most strongly upregulated proteins with dose-dependent patterns in *B. schlosseri* are many proteins known to be sensitive stress bioindicators in other invertebrates (Supplementary Table S1). Putative beta-lactamase-like protein 1 was among the most highly modulated proteins, showing a 10.4-fold increase at C3 and a 19.9-fold increase at C4. Mucin-5B also exhibited substantial up-regulation, with fold changes of 7.9 and 13.9 at C3 and C4, respectively. Body wall muscle protein HR-29 rose by 4.7-fold at C3 and 8.9-fold at C4, while universal stress protein Slr1101 increased 2.4-fold and 3.3-fold at C3 and C4, respectively. These expression patterns suggest clear toxic effects of copper on the tunicate proteome, which intensify in a dose-dependent manner.

**Figure 3.**
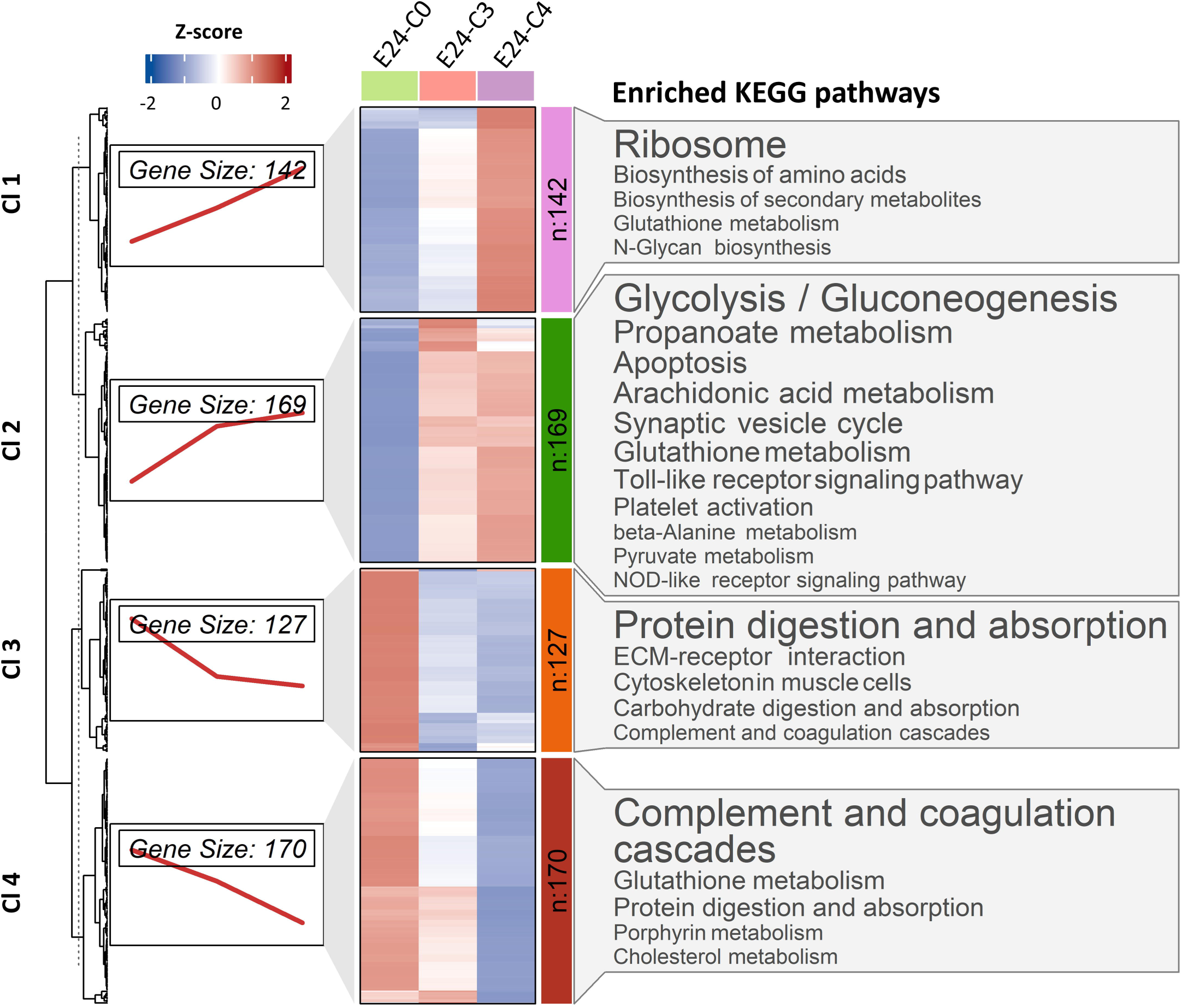
**Clustering and functional analysis of *B. schlosseri* proteins modulated by copper**. Differentially abundant proteins (DAPs) were clustered into four groups (Cl 1–4) based on expression trends across increasing copper concentrations. Line plots on the left show cluster size and expression dynamics. The heatmap presents standardized expression values (Z-scores) for each sample. Enriched KEGG pathways associated with each cluster are shown on the right, color-coded by functional category. Pathway label size is inversely proportional to p-value, with larger labels indicating stronger statistical significance.

To explore the biological functions related to these copper-induced proteome regulatory patterns, KEGG pathway enrichment analysis was conducted for each cluster (Figure 3). Glutathione metabolism, a key pathway involved in oxidative stress response, was enriched in Clusters 1, 2, and 4, with a higher number of DAPs observed in Cluster 2. In Cluster 1, the ribosome pathway was the most significant enriched pathway, followed by the biosynthesis of amino acids, pointing to enhanced protein synthesis. Cluster 2 showed strong enrichment in gluconeogenesis and propanoate metabolism, indicating a metabolic shift to support energy homeostasis under copper-induced stress. Arachidonic acid metabolism, involving several cytochrome P450 proteins, and apoptosis were also significantly enriched in Cluster 2. Additional pathways, including NOD-like and Toll-like receptor signaling and the synaptic vesicle cycle, point to alterations in innate immune responses and potential impacts on the nervous system. Clusters 3 and 4, composed of down-regulated DAPs, were mainly enriched in pathways related to the complement and coagulation cascades as well as protein digestion and absorption, suggesting a suppression of immune function and a potential decline in nutrient processing. While complement and coagulation cascades were more significantly enriched in Cluster 4, protein digestion and absorption pathways were more evident in Cluster 3. Cluster 3 also showed enrichment of the ECM–receptor interaction pathway, revealing a possible structural disruption of the extracellular matrix that may compromise tissue integrity and impair cellular communication under copper stress.

#### 3.2.2. Protein dynamics during recovery

To investigate the molecular basis of recovery from copper-induced stress, we compared the proteomes of tunicates sampled after 24 hours of recovery (R24) with those collected immediately after 24 hours of copper exposure (E24). A total of 2,618 differentially abundant proteins (DAPs) were identified across all conditions (C0, C3 and C4). Among them, 1,463 DAPs (55%) were identified in the control group (C0), likely reflecting physiological changes associated with the normal blastogenic cycle. To isolate recovery-specific signatures and minimize confounding effects of blastogenesis, DAPs identified in C0 were excluded from the analysis to keep the focus on proteins specifically upregulated or downregulated in the C3 and C4 recovery groups.

Among the upregulated DAPs (Figure 4A), the Upset analysis revealed that only 14 DAPs were common to all copper concentrations, highlighting distinct molecular signatures between recovery conditions from mild versus severe copper stress. A total of 373 upregulated DAPs were modulated in both C3 and the control condition (C0), but not in C4. In contrast, C4 shared only 4 DAPs with C0 that were not modulated in C3. Notably, 417 and 192 DAPs were uniquely upregulated in C3 and C4, respectively, with an additional 94 DAPs shared between the two conditions but absent in C0 controls. These subsets of protein groups, not modulated in C0, were selected for downstream functional enrichment analysis (Figure 4B). The functional analysis showed that most of the KEGG pathways enriched in C4 were also enriched in C3 but involved a greater number of DAPs in C4. This is particularly evident for immune-related KEGG pathways such as lysosome, phagosome, and complement cascades, pointing to an activation of proteolytic and immunity processes during recovery. Several pathways were mainly or exclusively enriched in recovery condition C3 indicating that full recovery processes take longer to initiate after more severe copper stress (C4). Among them, those related to extracellular matrix (ECM) interactions, focal adhesion, and cell adhesion molecules were the most significant enriched pathways, indicative of dynamic tissue remodeling. Importantly, all integrins (11) and most of the Laminin subunit proteins, which are associated with the PI3K-Akt signaling pathway, focal adhesion, and ECM-receptor interaction, were significantly upregulated exclusively in C3 (Supplementary Table S1), highlighting a strong activation of integrin-signaling pathways. C3 recovery also displayed a broad increase of intracellular signaling pathways such as MAPK, Ras, Rap1, Apelin, and cGMP-PKG, which are commonly associated with cell survival, proliferation, and stress adaptation. Interestingly, neuroactive ligand–receptor interaction and synaptic signaling were also exclusive to C3, potentially reflecting biological changes in the nervous system. Other pathways that were exclusively upregulated during C3 recovery included metabolic pathways related to protein digestion, carbohydrate metabolism, and sphingolipid metabolism.

**Figure 4.**
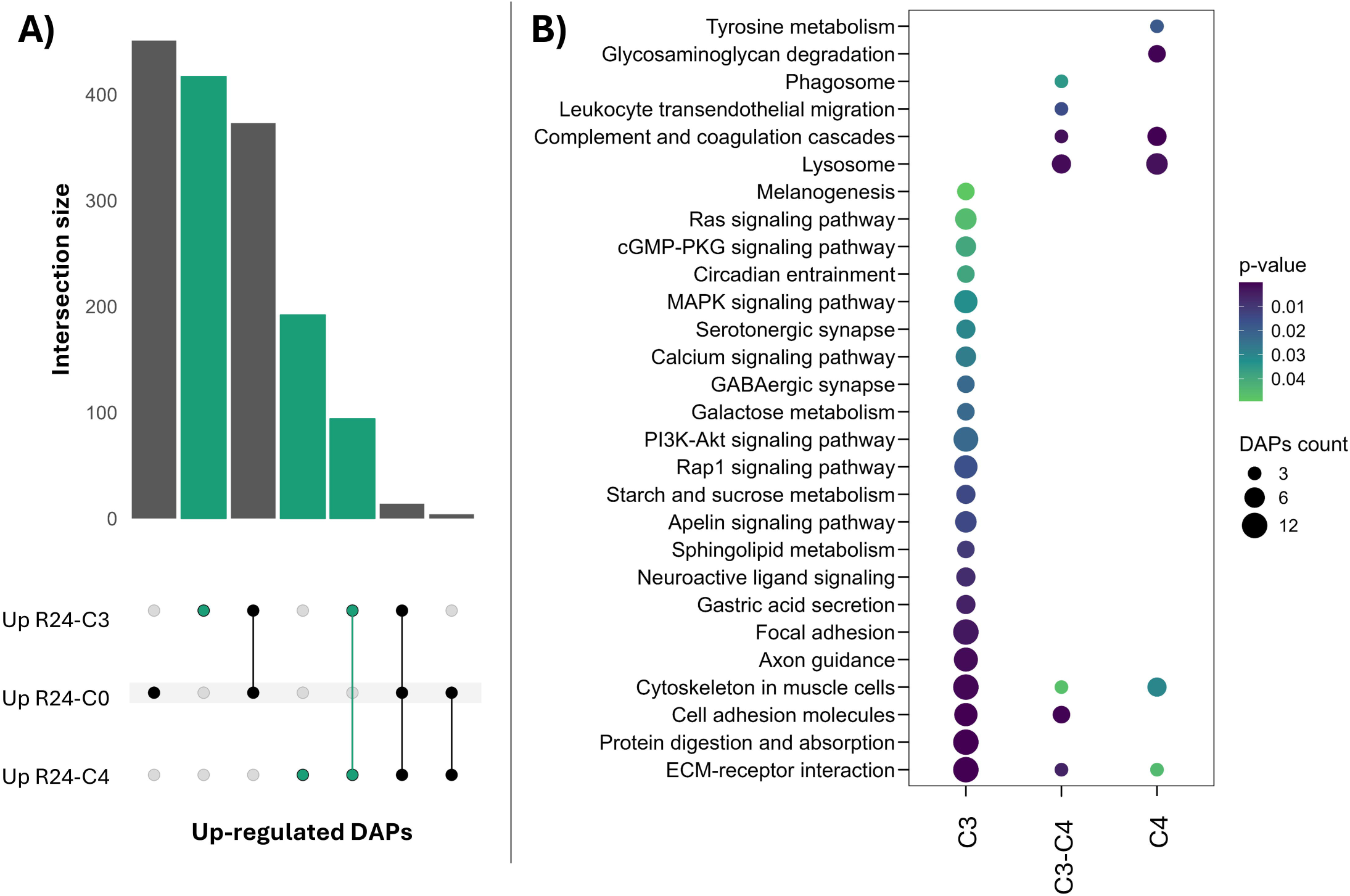
Functional analysis of up-regulated proteins after 24 hours of recovery from copper exposure. (A) Upset plot showing the number of differentially abundant proteins (DAPs) that are uniquely or commonly up-regulated across the different experimental conditions. Connected dots mean shared DAPs between those conditions, while single dots show DAPs unique to one condition. Bars highlighted in green indicate proteins up-regulated during recovery from copper exposure, but not in the control condition. These proteins were selected for further functional analysis. (B) Bubble plot showing KEGG pathways significantly enriched among up-regulated DAPs that are either specific to or shared between the C3 and C4 recovery conditions.

A similar trend is observed for the downregulated proteins during recovery (Figure 5). Upset plot analysis revealed that only 15 DAPs decreased across all recovery conditions (Figure 5A). A substantial overlap was observed between C0 and C3, with 387 shared DAPs, whereas C4 shared only 20 DAPs with the control condition. 371 and 304 DAPs decreased in C3 and C4, respectively, while 72 DAPs decreased in both recovery conditions. DAPs specifically downregulated in C3 and C4 or in both conditions were further analyzed for functional enrichment (Figure 5B). Downregulated DAPs from both C3 and C4 conditions were strongly enriched in KEGG pathways related to primary metabolism, indicating substantial metabolic reprogramming during the recovery phase. This included coordinated downregulation of pathways involved in carbohydrate, lipid, and amino acid metabolism. Detoxification-related pathways, such as peroxisome function and glutathione metabolism, were also suppressed, pointing to the decline in oxidative stress as recovery progressed. Only a few pathways were uniquely downregulated in C3, including the HIF-1 signaling pathway, which is linked with immune system. More KEGG pathways were exclusive to C4. Most of them were related to digestive functions, including significant suppression of protein digestion, gastric acid secretion, and pancreatic secretion pathways. Additional specific enriched pathways included TCA cycle and other core metabolic processes.

**Figure 5.**
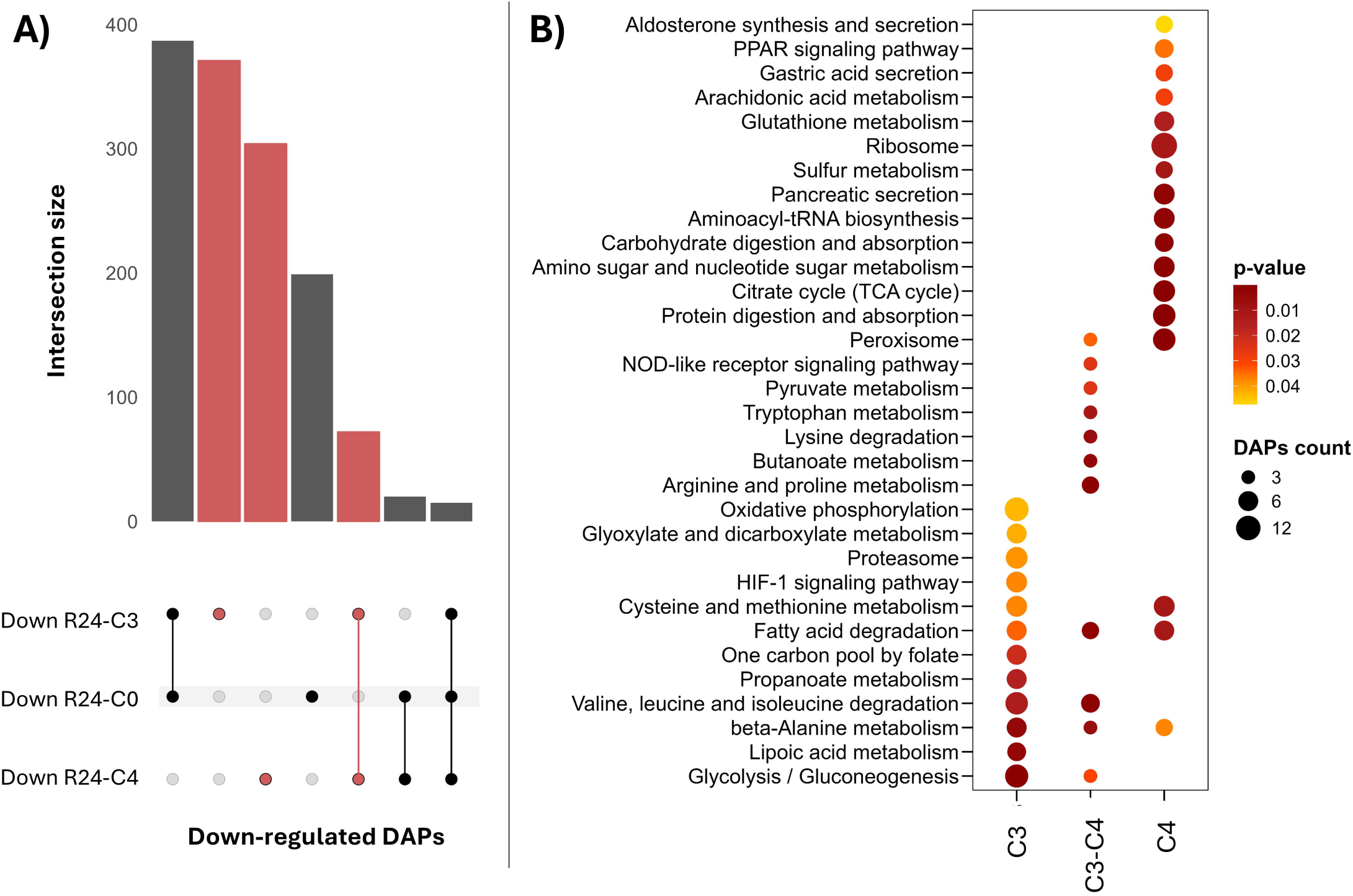
Functional analysis of down-regulated proteins after 24 hours of recovery from copper exposure. (A) Upset plot showing differentially abundant proteins (DAPs) that are uniquely or commonly up-regulated across the different experimental conditions. Bars highlighted in red indicate proteins down-regulated during recovery from copper exposure, but not in the control condition. These proteins were selected for further functional analysis. (B) Bubble plot showing KEGG pathways significantly enriched among down-regulated DAPs that are either specific to or shared between the C3 and C4 recovery conditions.

#### 3.2.3. Molecular signatures of phenotypic changes

To identify reliable molecular signatures of copper stress, we targeted DAPs whose expression patterns closely reflected the phenotypic changes observed during exposure and recovery to copper. These alterations included reduced body area and darker pigmentation during exposure to C3 and C4 copper concentrations, followed by increased body area and lighter pigmentation during recovery, which was evident in C3 but absent in C4. To this end, the expression dynamics of the 223 DAPs identified after 24 hours of exposure to C3 were analyzed over time in both C3 and C4 conditions using clustering analysis (Supplementary Figure S3). This strategy provided the opportunity to reveal two distinct signatures of proteins with contrasting expression trajectories. Signature 1 included 13 DAPs that were up-regulated in C3 during exposure, returned to lower levels during recovery, but remained consistently high in C4 across both phases. Signature 2 comprised 25 DAPs that were down regulated in C3 during exposure but returned to baseline levels during recovery, while remaining suppressed in C4. After excluding redundant functional annotations, a final set of 35 distinct DAPs was retained, representing robust protein signatures associated with contrasting stress and recovery dynamics (Figure 6).

**Figure 6.**
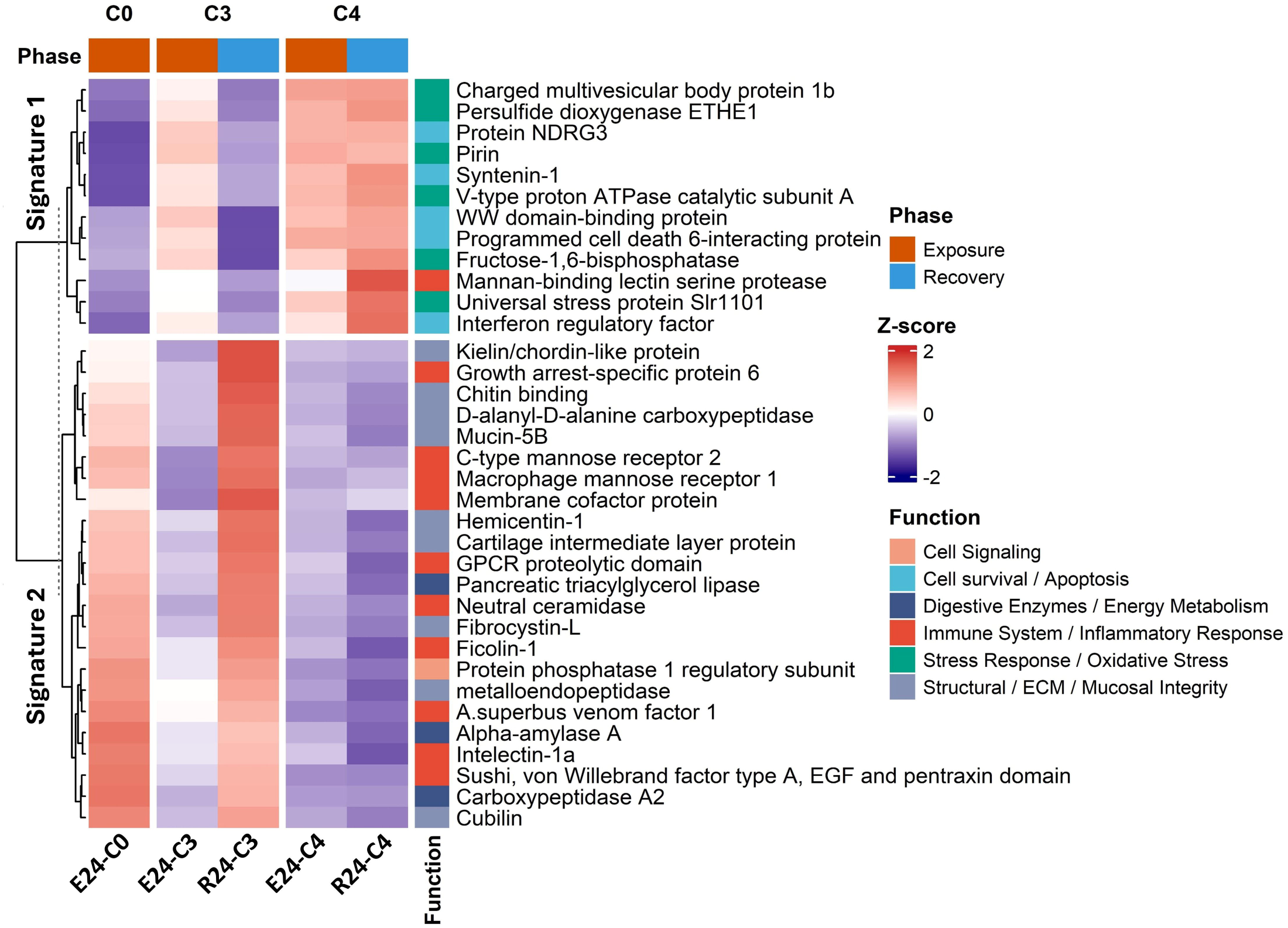
Expression patterns and functional annotation of candidate molecular markers across copper exposure and recovery. Heatmap showing the Z-score-normalized expression of selected DAPs grouped by clustering into Signature 1 and Signature 2 across C0, C3, and C4 conditions. Protein expression dynamics are shown for both exposure (E) and recovery (R) phases. Functional categories highlight involvement in cell signaling, oxidative stress, apoptosis, immune response, and structural integrity, supporting their relevance as stress and recovery markers.

To gain functional insight into these molecular signatures that mirror whole organismal phenotypes observed in copper-exposed *B. schlosseri*, the corresponding 35 DAPs were manually categorized based on their annotated biological functions (Figure 6). DAPs comprising signature 1 were primarily associated with immune and inflammatory responses, cell survival and apoptosis, and responses to cellular or oxidative stress. For example, the universal stress protein Slr1101 increased more than twofold during exposure in both C3 and C4, returned to baseline during recovery in C3, but remained elevated in C4. Similarly, Pirin, known as a sensor of oxidative stress, showed a comparable pattern of modulation across conditions. Programmed cell death interacting protein was also elevated during exposure, followed by down-regulation during recovery in C3. DAPs comprising signature 2 were predominantly involved in digestive processes and energy metabolism, immune function, and related to extracellular matrix organization and mucosal integrity. For example, alpha-amylase and mucin-5B both decreased significantly during exposure and returned to baseline during recovery in C3, while remaining low in C4. Mannose-binding lectin receptor, which participates in pathogen recognition and immune signaling, followed a similar trend.

## 4. Discussion

This study provides an integrative assessment of the stress and recovery dynamics in the colonial tunicate *B. schlosseri* following acute copper exposure. By combining whole organism phenotypic measurements with quantitative proteomics, we identify coordinated changes at both the organismal and molecular levels that reflect the organism’s sensitivity and ability to recover from exposure to this environmental contaminant.

Among the phenotypic markers, changes in tunicate system area and pigmentation emerged as robust and quantifiable indicators of copper-induced stress. These responses were clearly dose-dependent, with higher copper concentrations leading to more pronounced reductions in system area and increased pigmentation. This pattern enabled the detection of sublethal effects and helped identify the lowest concentration at which copper began to impair tunicate physiology. Moreover, these markers changed during recovery, allowing to differentiate *B. schlosseri’s* recovery capacity across copper levels and identify threshold concentrations causing lasting effects. These traits provide a more reliable and objective assessment compared to traditional botrylliod tunicate physiology markers, which are mostly qualitative and based on visual observations such as ampullae health, siphon opening, and general colony appearance (Gregorin et al., 2021). While these observations successfully identified toxic copper concentrations in *B. schlosseri*, they remain subjective and prone to observer bias. This caveat makes it hard to monitor stress consistently, especially in the early stages of stress exposure. A few quantitative markers have also been explored, such as heartbeat and colony growth rates (Dijkstra et al., 2008; Osborne et al., 2018). However, monitoring heart movements is technically challenging and often hindered by pigmentation that obscures visualization. Colony growth and area were effective markers for comparing how ascidian species tolerate copper exposure (Osborne et al., 2018), however, these markers are mainly suited for chronic studies. In this study, changes in system area and pigmentation provided a sensitive and practical means to assess both copper-induced stress and recovery capacity. These quantitative markers guided the selection of relevant copper concentrations for subsequent molecular investigations through proteomic analysis.

Proteomic analyses were performed on *B. schlosseri* colonies exposed to moderate (300 µg/L, C3) and high (900 µg/L, C4) copper concentrations, which caused significant phenotypic changes, with only the colonies exposed to the moderate concentration (C3) showing signs of recovery. In line with the organismal phenotype responses, quantitative proteomics of exposed-tunicates revealed a dose-dependent modulation of protein abundances, with stronger changes of molecular phenotype at higher copper concentration. The most strongly up-regulated proteins in response to copper exposure included several stress-related factors, such as Universal stress protein Slr1101, Mucin-5B and the body wall muscle protein HR-29. Universal stress proteins are a highly conserved family of proteins found across bacteria, archaea, plants, and some invertebrates and act as central regulators of stress responses, supporting growth, development, and resilience under diverse environmental stresses (Forêt et al., 2011; Vollmer and Bark, 2018). In invertebrates, mucins have been increasingly reported for their involvement in stress responses and as ingredients of mechanical and chemical mucus barriers that afford protection against pathogens and environmental toxins (Mentino et al., 2025). The chaperone protein HR-29, identified as a biomarker of soft tissue disorders in the tunicate *Halocynthia roretzi*, is associated with immune responses and the maintenance of structural integrity (Cha et al., 2011).

Further functional analyses of the proteome revealed enrichment of key stress response pathways. These included glutathione metabolism, a well-established defense against oxidative damage (Kültz, 2020; Raj Rai et al., 2021), and arachidonic acid metabolism, which contributes to oxidative signaling, inflammation, and cellular proliferation (Kültz, 2020; Tallima and El Ridi, 2018). Glutathione-related proteins have been shown to respond to copper in various invertebrates, including ascidians (Li et al., 2009; Franchi et al., 2012; Peng et al., 2024), while arachidonic acid derivatives are thought to be central mediators of copper tolerance, as reported in earthworms (*Eisenia fetida*) (Zhang et al., 2024). Conversely, down-regulated proteins were mainly associated with immune and digestive functions. This suppression aligns with the known immunotoxic and digestive-disrupting effects of heavy metals, including copper, and general suppression of vegetative bodily functions as a non-specific response to many types of stress (Selye, 1936). For example, copper exposure in mussels has been shown to affect immune parameters such as hemocyte abundance, phagocytic activity, and complement system components (Chen et al., 2018; Pipe et al., 1999). In *Gammarus fossarum*, metal exposure led to dose-dependent reductions in feeding activity and digestive protein expression (Leprêtre et al., 2023) and similar inhibition of digestive enzyme activity has been reported in other marine invertebrates (Chen et al., 2002). Altogether, proteome changes induced by copper exposure shed light on the trade-offs and conserved mechanisms to metal-induced stress. They reflect a coordinated molecular stress response, characterized by the upregulation of proteins involved in detoxification and oxidative signaling, alongside the downregulation of immune and digestive mechanisms.

The contrasting recovery trajectories between moderate and high copper exposure offered further insights into the molecular basis of regeneration. In colonies exposed to both moderate and high copper concentrations, early recovery was marked by metabolic downregulation, particularly in carbohydrate, lipid, and amino acid pathways, suggesting a shift towards repair and survival processes (Eming et al., 2017; Kültz, 2005). This was accompanied by the downregulation of glutathione and peroxisome pathways, indicating a reduction in oxidative stress as recovery progressed. In recovering colonies, upregulated pathways were strongly associated with cell proliferation and angiogenesis signaling (e.g., PI3K, Apelin), alongside components of the extracellular matrix (ECM), particularly those involved in cell adhesion. Notably, the upregulation of all integrins highlighted the critical role of ECM-receptor interactions in regeneration (Cheong et al., 2025). These molecules help maintain tissue integrity, mediate cell– matrix interactions, and activate downstream pathways like PI3K-AKT, which regulate cell survival, proliferation, and migration (Mezu-Ndubuisi and Maheshwari, 2021). The enrichment of ECM-receptor interaction and integrin signaling pathways aligns with previous studies on whole-body regeneration (WBR) in botrylloid tunicate species (Kassmer et al., 2020; Ricci et al., 2022). These findings suggest that ECM components, angiogenic factors, and integrins are central to the regenerative processes that support recovery after injury and chemical stress, further underscoring their role in tissue repair and stem cell activation. Interestingly, several upregulated pathways were linked to the nervous system, suggesting that copper exposure impacts nervous system function, further supported by the enrichment of nervous system-related pathways during the exposure phase. Previous research has shown that copper toxicity impairs neural function, behavior, and development in various invertebrate species (Brown et al., 2004; Hwang et al., 2014; Zhang et al., 2021). *B. schlosseri* exposed to higher copper levels (C4) failed to recover and showed persistent suppression of energy metabolism and digestive function. Copper toxicity at high concentrations may overwhelm cellular processes, impair essential functions, and surpass the organism’s capacity for recovery. This difference between proteome dynamics during recovery from moderate (C3) versus severe (C4) copper stress highlights the critical role of energy metabolism and structural remodeling in post-exposure stress recovery.

Finally, the identification of molecular signatures that mirror phenotypic changes in *B. schlosseri* offers a powerful framework for understanding organismal stress responses and provides key entry points for future applications in stress biology and evolutionary research. These signatures include proteins associated with oxidative stress responses, immune modulation, apoptosis, energy metabolism, and tissue integrity. They are indicative of biological processes that are broadly mobilized in response to environmental stress-induced macromolecular damage across a wide range of animal taxa. Their consistent association with system pigmentation and size changes enhances their relevance as integrated physiological markers of stress severity. Moreover, the specific proteomic signatures associated with copper stress, even though comprised of many non-specific stress proteins, may entail specific patterns that are unique to copper stress relative to other types of environmental stress when comparing topological information inherent in these proteomic signatures. Future experiments utilizing other types of stress in a comparable experimental design can address this hypothesis. Beyond mechanistic insight, these molecular signatures can serve as sensitive tools in ecotoxicology, enabling early detection of sublethal stress and expanding the repertoire of bioindicators in *B. schlosseri*, where few molecular indicators exist. Furthermore, reliable signatures of stress-induced macromolecular damage (including mutagenesis) and phenotypic plasticity on the one hand and rapid recovery on the other hand could serve to optimize stress-induced evolution experimental designs aimed at maximizing adaptive mutagenesis and artificial selection of tunicates that are capable of asexual whole body regeneration.

## 5. Conclusion

In conclusion, this study provides a comprehensive understanding of copper-induced stress and recovery mechanisms in *B. schlosseri*, combining whole organism phenotyping and quantitative proteomics. The identification of robust bioindicators of copper stress, such as system area reduction and pigmentation darkening, alongside specific molecular signatures identified in the study offer valuable tools for environmental monitoring and risk assessment. Proteomic analyses further clarified the molecular basis of copper toxicity, revealing specific proteins involved in conserved stress response mechanisms such as oxidative defense, immune modulation, and metabolic reprogramming. In addition, the recovery phase was marked by the upregulation of extracellular matrix components, adhesion molecules, and proliferative signaling pathways, highlighting the most critical proteins for active tissue repair processes and the organism’s capacity for regeneration following toxic exposure.

Beyond its ecotoxicological relevance, this work contributes significantly to future research on stress-induced evolution (SIE). The use of sublethal copper concentrations (e.g., 300 µg/L) that induce physiological and molecular responses indicative of increased mutagenic damage without causing mortality offers a relevant framework for modeling adaptive stress responses over time. Such intermediate stress levels generate the selective pressure necessary to drive evolutionary processes while preserving organismal viability and regenerative capacity. Crucially, the quantitative proteomics approach presented here lays the molecular foundation for investigating how environmental stress shapes evolutionary trajectories, by identifying stable, quantifiable molecular signatures associated with specific stress exposure and recovery regimens.

## CRediT authorship contribution statement

### Maxime Leprêtre

Conceptualization, Methodology, Formal analysis, Writing – original draft. **Dietmar Kültz:** Supervision, Conceptualization, Methodology, Funding, Writing – review & editing.

## Data availability

All raw proteomics data, metadata, and DIA-NN analysis results are available publicly in the PRIDE Archive (accession: PXD064899) via the ProteomeXchange Consortium (Perez-Riverol et al., 2025).

## Acknowledgement

This work was supported by NSF grant MCB-2127516.

## Supplementary Files

**Supplementary Figure S1.** Visual Comparison of Tunicate Responses to Copper Exposure and Recovery. Representative images of G2 (A) and G3 (B) tunicate systems at baseline (BL), after 24 h of copper exposure (E24), and during recovery at 24 h (R24) and 96 h (R96), under three copper concentrations: C0 (0 µg/L), C3 (300 µg/L), and C4 (900 µg/L). All images were taken at the same magnification and light intensity to allow direct visual comparison.

**Supplementary Figure S2.** Dose-dependent effects of copper exposure on system pigmentation and recovery across the three genotypes. (A) Boxplots showing the percentage change in system pigmentation relative to baseline (BL) after 24 hours of exposure (E24) across five copper concentrations (C0 = 0 µg/L, C1 = 30 µg/L, C2 = 100 µg/L, C3 = 300 µg/L, C4 = 900 µg/L) in genotypes G1, G2, and G3. Different letters indicate statistically significant differences between conditions within each genotype (Tukey’s HSD, p < 0.05). (B) Line plots showing the dynamics of system pigmentation change over time, during exposure (BL–E24) and recovery (R24–R96) for each genotype. Line colors correspond to copper concentrations and points represent mean values ± standard error. Dashed vertical line marks the end of exposure and beginning of the recovery.

**Supplementary Figure S3.** Distinct protein expression trajectories reflecting phenotypic recovery dynamics. K-means clustering of the 223 DAPs identified after C3 exposure revealed two main expression groups across C3 and C4 conditions. Group 1 includes proteins upregulated during C3 exposure that returned to baseline during recovery but remained elevated in C4. Group 2 comprises proteins downregulated during C3 exposure that recovered only in C3. Venn diagrams highlight the overlap of proteins showing consistent trends across both conditions.

**Supplementary Table S1.** Functional annotation and differential proteomic analysis of proteins in *Botryllus schlosseri* exposed to and recovered from varying copper concentrations.

