## Supplementary figures and images for "Copper-Induced Stress and Recovery Impacts on Organismal Phenotypes and the Underlying Proteomic Signatures in *Botryllus schlosseri*"

### Supplementary Figure S1

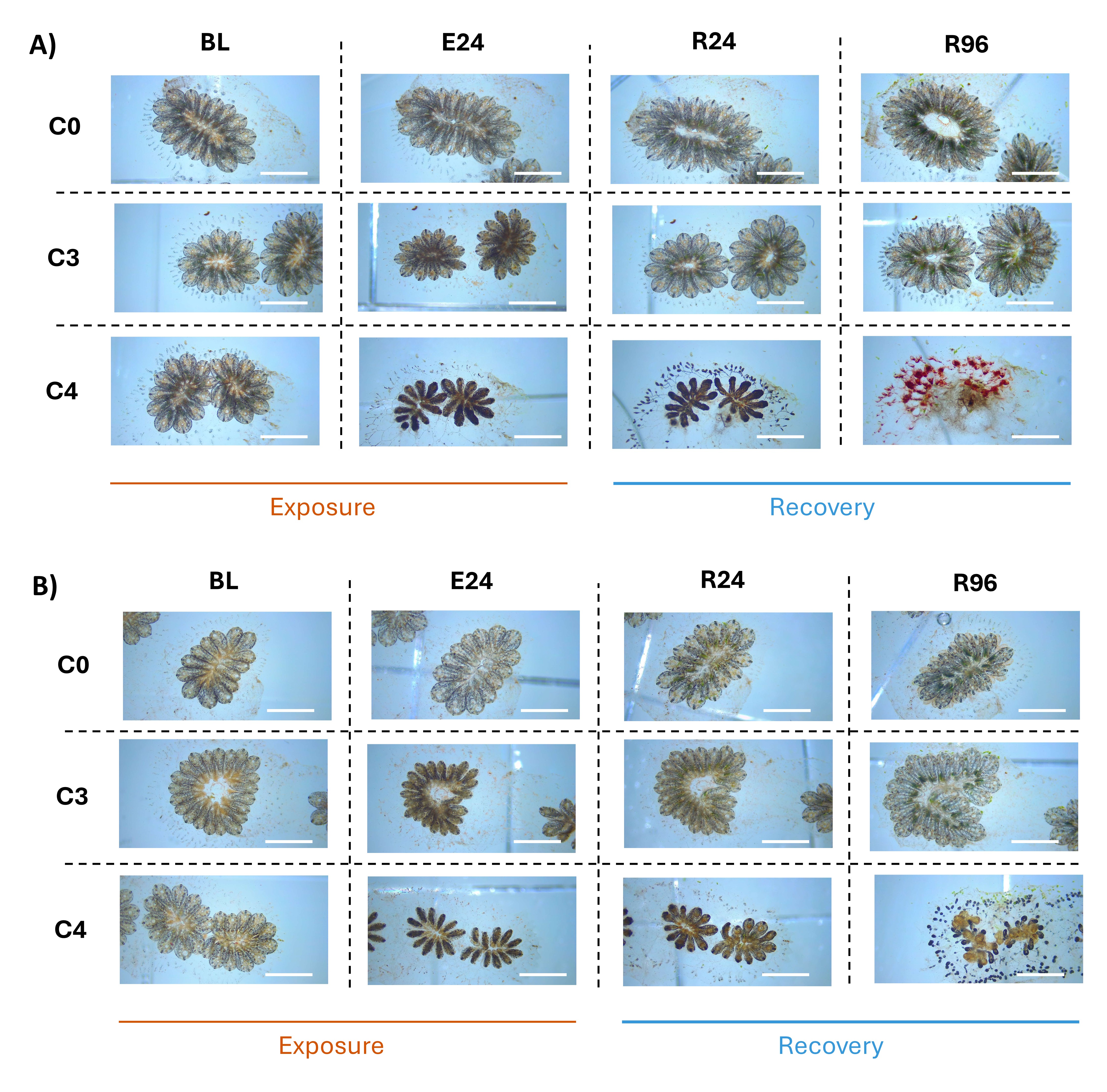

### Supplementary Figure S2

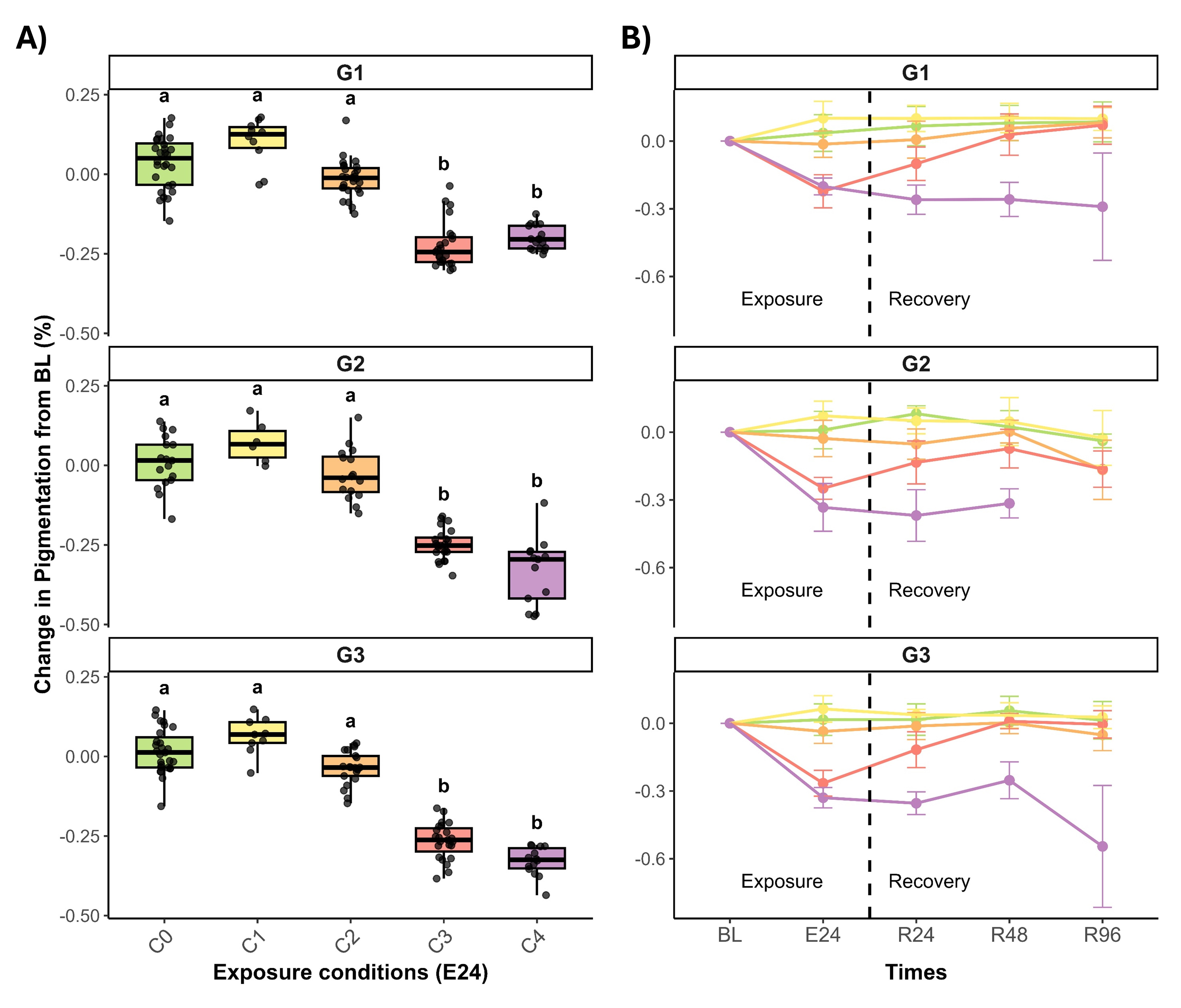

### Supplementary Figure S3

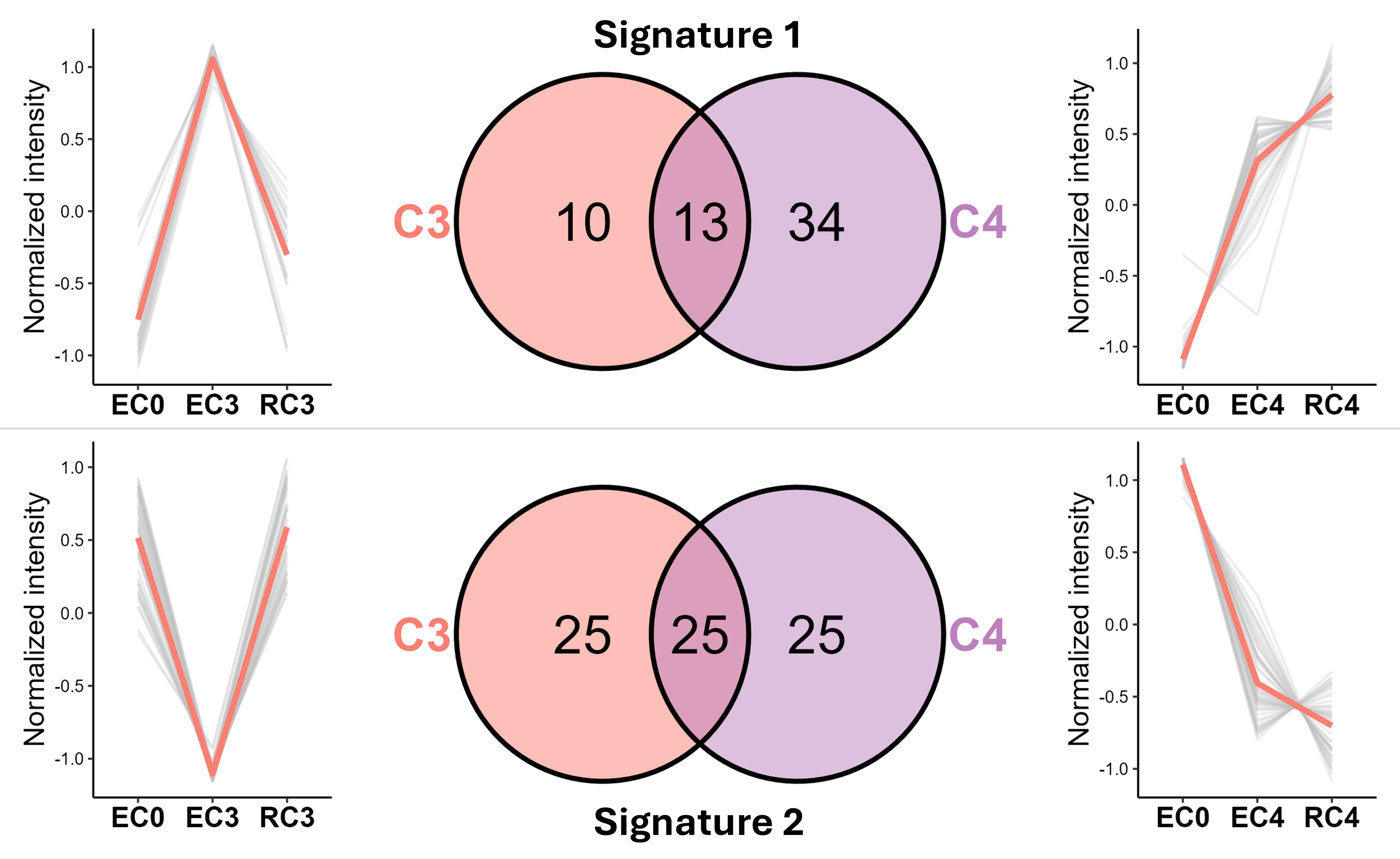
